# Dynamics of visual contextual interactions is altered in Parkinson’s disease

**DOI:** 10.1101/549691

**Authors:** M. Isabel Vanegas, Annabelle Blangero, James E Galvin, Alessandro Di Rocco, Angelo Quartarone, M. Felice Ghilardi, Simon P. Kelly

**Affiliations:** Department of Biomedical Engineering, The City College of The City University of New York, New York, New York, USA; Department of Ophthalmology and Visual Sciences, University of Utah, Salt Lake City, Utah, USA; OCTO Technology, Paris, France; Charles E. Schmidt College of Medicine, Florida Atlantic University, Boca Raton, FL; Parkinson’s and Movement Disorders, Northwell Health, New York, NY; IRCCS Centro Neurolesi Bonino Pulejo, Department of Biomedical, Dental Science and Morphological and Functional Images, University of Messina, Messina, Italy; Department of Physiology, Pharmacology & Neuroscience, CUNY School of Medicine, New York, New York, USA; School of Electrical and Electronic Engineering, University College Dublin, Belfield, Dublin 4, Ireland

**Keywords:** Parkinson’s disease, EEG, ssVEP, surround suppression, contrast response function, adaptation, gain control

## Abstract

Over the last decades, psychophysical and electrophysiological studies in patients and animal models of Parkinson’s disease (PD), have consistently revealed a number of visual abnormalities. In particular, specific alterations of contrast sensitivity curves, electroretinogram (ERG), and visual evoked potentials (VEP), have been attributed to dopaminergic retinal depletion. However, fundamental mechanisms of cortical visual processing, such as normalization or “gain-control” computations, have not yet been examined in PD patients. Here we measured electrophysiological indices of gain control in both space (surround suppression) and time (sensory adaptation) in PD patients based on steady-state VEP (ssVEP). Compared to controls, patients exhibited a significantly higher initial ssVEP amplitude that quickly decayed over time, and greater relative suppression of ssVEP amplitude as a function of surrounding stimulus contrast. Meanwhile, EEG frequency spectra were broadly elevated in patients relative to controls. Thus, contrary to what might be expected given the reduced contrast sensitivity often reported in PD, visual neural responses are not weaker; rather, they are initially larger but undergo an exaggerated degree of spatial and temporal gain control and are embedded within a greater background noise level. We conclude that compensatory cortical mechanisms may play a role in determining dysfunctional center-surround interactions at the retinal level.

## Introduction

Parkinson’s disease (PD) is a progressive neurological disorder characterized by dopamine deficiency most famously in striatal circuits of the basal ganglia, but also in other dopamine-regulated systems, including the retina.^1-3^ The involvement of circuits other than the basal ganglia could explain some of the non-motor symptoms including sleep regulation problems, autonomic dysfunction, hyposmia and visual abnormalities.^4^

Visual symptoms in PD range from blurry vision, decreased ability to discern color, loss of contrast sensitivity to circadian dysregulation and visual hallucinations.^5-7^ Systematic abnormalities have also been reported by electrophysiological and psychophysical studies. Pattern electroretinogram (PERG) responses in patients and animal models are decreased in amplitude and increased in latency compared to controls^8-12^, particularly in an intermediate range of spatial frequencies (2-5 cpd) where contrast sensitivity is highest.^9^,^11^ Pattern Visual-evoked potential (PVEP) studies have shown parallel effects^13-15^, including delayed latency of the N70 and P100 component in a similar range of spatial frequencies and for temporal frequencies between 4-5 Hz^16^, and phase delay of a low-frequency steady state response.^17^ Acute levodopa administration in general reversed both PERG and PVEP abnormalities, suggesting that dopamine has an important role in visual processing.^11-15^,^17^ Psychophysical studies of spatiotemporal tuning curves in PD patients have confirmed a major loss of contrast sensitivity for spatial frequencies in the 2-4 cpd range and temporal frequencies between 4 and 8Hz.^18-20^ Given the overlap in the affected spatial and temporal frequency ranges, it is plausible that these perceptual abnormalities are linked to the retinal dysfunction reflected in the PERG, and both may arise from the dopaminergic cell degeneration that has been observed in the retina in PD.^2^,^21-23^ Dopaminergic amacrine cells are present in the retinal interplexiform layer and are key players in neural transmission from photoreceptors to ganglion cells and in the generation of center-surround interactions of ganglion cell’s receptive fields, thus contributing to spatial frequency tuning.^23-25^ Further, mouse models of retinal dopamine deficiency have shown loss of contrast sensitivity and visual acuity, resembling the visual abnormalities described in PD.^26^

While those findings provide an extensive characterization of the retinally-mediated visual abnormalities present in PD, relatively little is known regarding potential differences in visual processing at the early cortical level, and, to our knowledge, there has yet been no systematic study of fundamental aspects of visual processing such as spatial and temporal gain control. Here we addressed this gap by examining cortically-generated, steady-state visual-evoked potentials (ssVEP) in PD and their modulation by surrounding spatial context and temporal adaptation.

We employed a recently developed paradigm designed to probe visual surround suppression whereby the perceived intensity of a stimulus decreases under the presence of a surrounding pattern^27,28^, a phenomenon that relies on lateral, feedback and feedforward connections from the earliest stages of information processing in the retina to visual cortex.^29,30^ Since the focus of our investigation was on cortical processing, we used spatial and temporal frequencies lying outside the ranges that typically reveal retinal abnormalities in PD (1cpd and 25Hz). We reasoned that, if the only alterations in cortex are those inherited from the retina, one should expect little or no difference in cortical responses.

We found several differences in patients with PD compared to healthy controls: 1) a greater background noise level in the frequency spectrum; 2) a higher visual response (ssVEP amplitude) that quickly adapts over time, and 3) a stronger relative suppression of the ssVEP amplitude under the presence of surrounding stimuli. These overall signatures of visual contextual interactions indicate that contrast gain control is abnormal in PD and thus may serve as biomarkers to aid diagnosis and to evaluate therapeutic efficacy.

## Results

Data were analyzed from a short surround-suppression paradigm recorded as part of a larger study of 28 PD patients and 30 healthy age-matched controls. In this paradigm, subjects passively viewed a series of 2.4-s long trials in which four “foreground” grating stimuli flickered on-and-off at 25Hz, embedded within a static (non-flickering) surround grating pattern, with both the foreground (FG) and surround (SS) contrasts varying randomly from trial to trial.

### Background (no-stimulus) Frequency spectrum profiles

Since differences in background spectral EEG amplitude have previously been reported in PD^31-34^ and would impact the estimation of ssVEP amplitudes driven by the external flicker-stimulation, it was important first to characterize such background spectral differences. EEG spectra computed from trials with zero foreground contrast and zero surround contrast (blank screen), averaged across a cluster of six parieto-occipital electrodes where ssVEPs were measured, revealed a broadly elevated spectrum in PD relative to controls (Figure 1). A bootstrap statistical test at the critical frequency of 25Hz revealed a significant difference in the means (p<0.05). Importantly, the spectral difference extended through the θ [4-7Hz], α [8-12Hz] and β [18-35Hz] bands.

**Figure 1.**
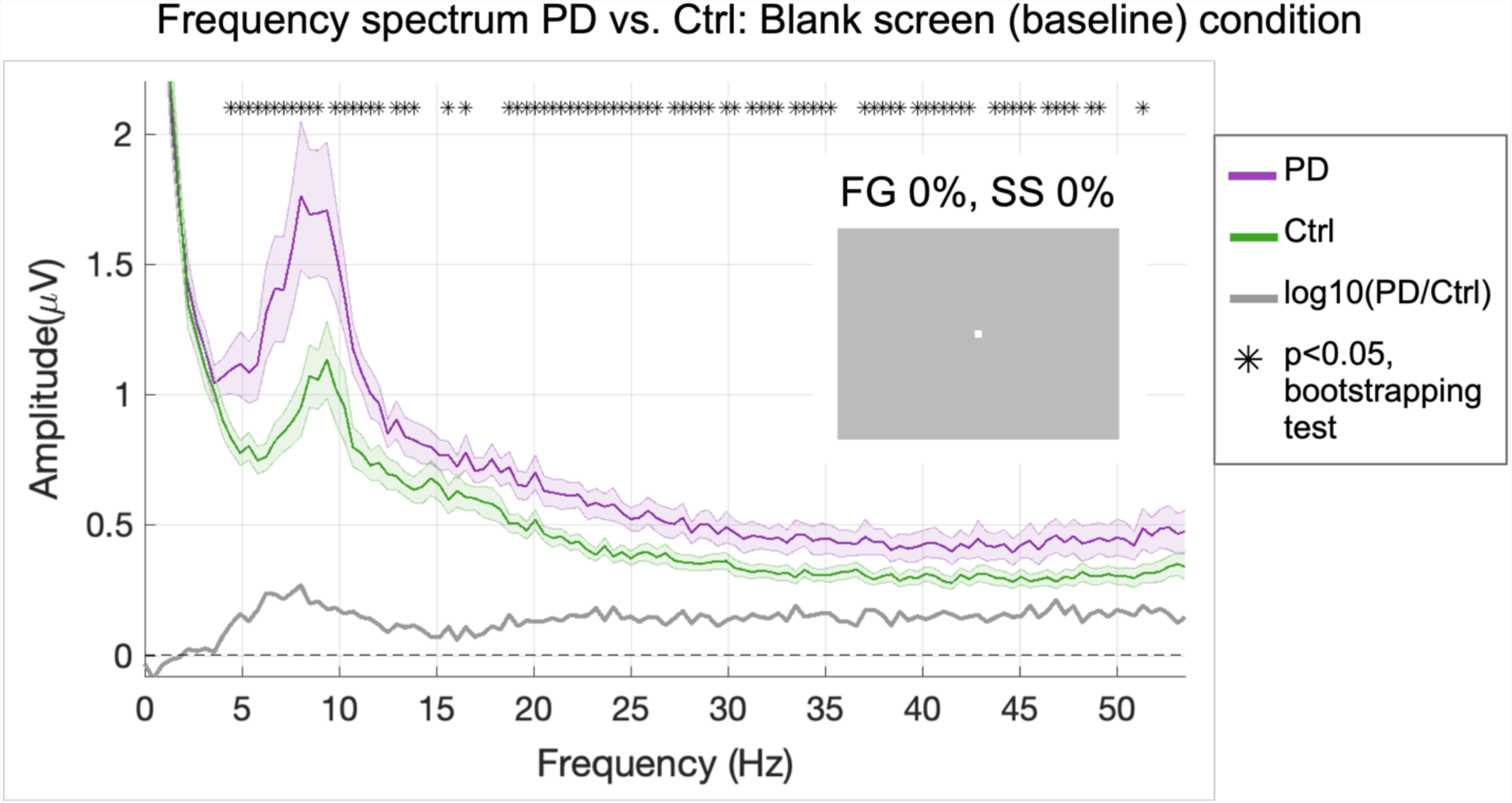
Background frequency spectrum profiles in patients versus controls. Fast Fourier Transforms were computed on the last 2.24 s of all trials with 0% foreground and 0% surround contrast (blank screen, shown in inset) and then averaged across trials and across subjects within each group. Patients showed significantly higher spectral amplitudes, marked with asterisks for each of the corresponding frequency bins (bootstrapping test, p<0.05). Shaded error bars indicate mean ± standard error of the mean (s.e.m.).

The broad spectral elevation seen under no stimulation conditions would also have the effect of elevating the amplitude of the 25-Hz ssVEP under the flicker-stimulus conditions when measured in the same way, even if there were no underlying differences in the amplitude of the exogenous response to the flicker. To eliminate this potential confound we took two measures. First, rather than averaging the single-trial spectra, we computed Fourier spectra on the average across trials per condition, thus averaging-out the background activity levels while retaining the ssvEP signal because it is strictly phase-locked to the stimulus (Figure 2A). Second, we further used the blank-screen condition to determine a lower number of trials to average within each control subject such that the background noise was elevated to the same average level as the PD patients. Tracing background noise levels in the 25Hz frequency bin during the no-flicker stimulus condition revealed that equal levels of background noise were achieved in both groups when the maximum number of available trials per condition was included for the PD group and about half of the total trials (3 to 4) for the controls (Figure 2B). This random trial rejection policy was then applied in the same way in the flicker conditions.

**Figure 2.**
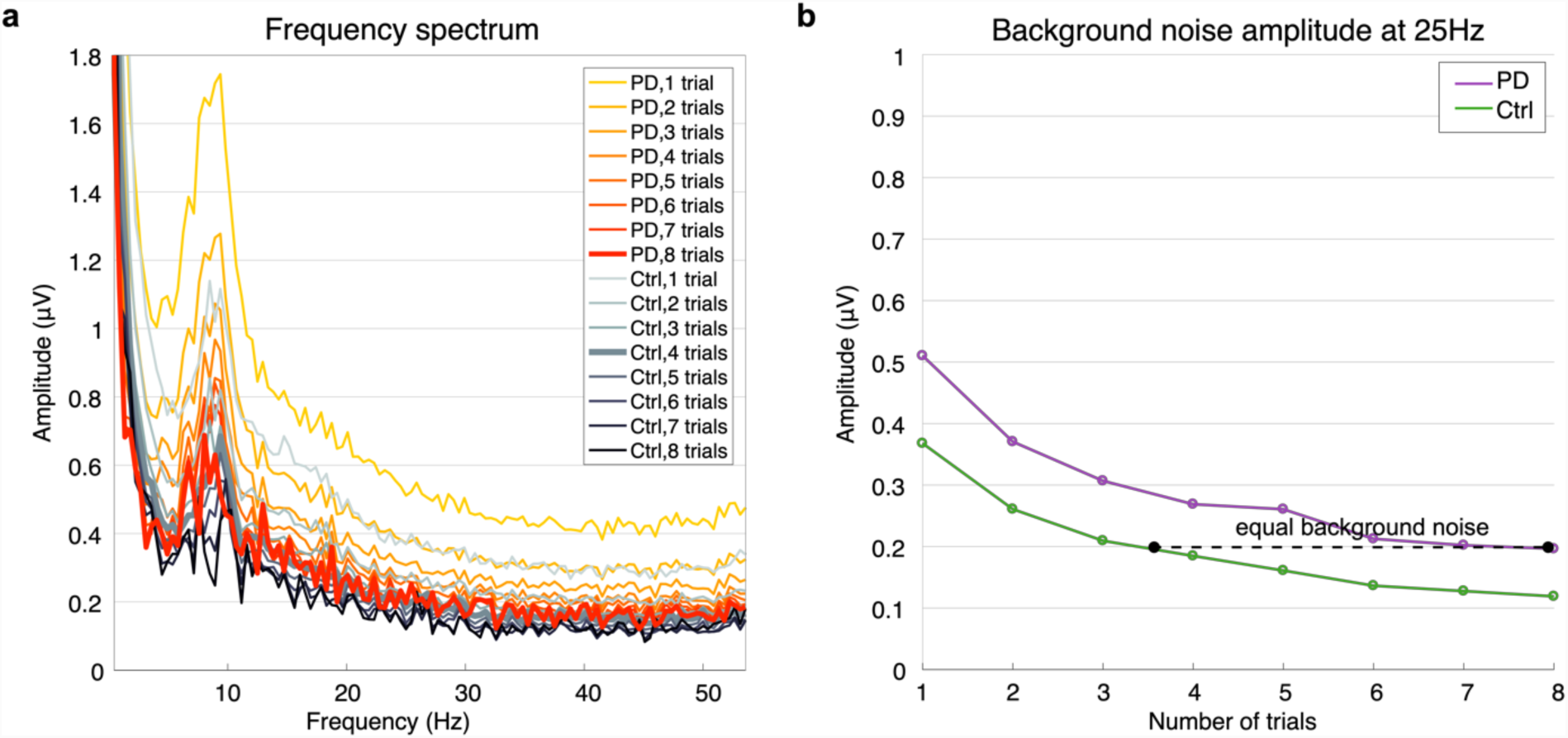
Background noise as a function of number of trials for both subject groups. **a-** Frequency spectrum for PD and Ctrl, computed using one to the maximum amount of trials in the blank-screen (baseline) condition of 0% foreground and 0% surround contrast. **b-**Amplitude at the 25Hz bin as a function of number of trials. A Fast Fourier Transform (FFT) was computed on the average time-domain signals across the given number of trials first, with that number selected randomly 1,000 times from the total trials available for that condition, and the resulting FFTs averaged. Background noise levels matched when the FFT was computed on the average signal across the maximum available trials (up to 8 per subject) for PD subjects and half of the total (up to 4 trials per subject) for control subjects.

### Surround suppression effects

Applying this noise equalization approach, we examined ssVEP contrast response functions and temporal profiles (Figures 3 and 4). These analyses were carried out on subsets of age-matched patients and controls without cognitive impairment (eleven per group, see methods). Contrast response functions show that ssVEP amplitude increased as a function of foreground contrast in both groups, and decreased as the contrast of the surround stimulus increased (Figure 3A,B).

**Figure 3.**
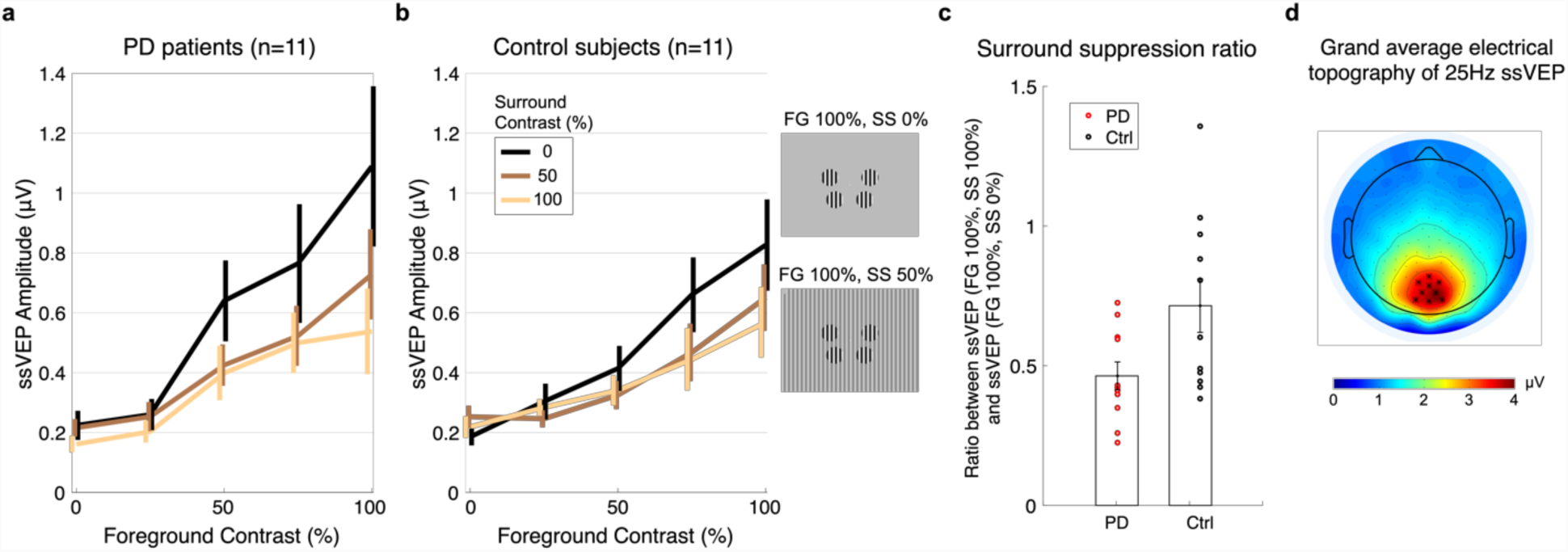
Contrast response functions for three different surround contrast conditions. **a-** Patients with Parkinson’s disease (PD). **b-**Control subjects. A fast Fourier transform (FFT) was computed for a 2,240-ms window beginning 160ms after stimulus onset to extract the single amplitude value at the frequency of stimulation 25Hz. Contrast response functions are constructed from the ssVEP amplitude in each stimulus configuration of foreground (FG) and surround (SS) contrasts. Foreground stimulus varied across five levels of contrast: 0, 25, 50, 75 and 100%. Surround stimulus varied across three levels of contrast: 0% contrast (black trace), 50% contrast (brown trace), and 100% contrast (yellow trace). Insets show stimulus configurations for FG 100% embedded in SS of 0% and 50% contrast. **c-**Surround suppression ratio. Each point corresponds to a subject. The suppression ratio is computed as the ratio between the ssVEP amplitude for FG=100%, SS=100% and the ssVEP amplitude for FG=100%, SS=0%. Error bars indicate s.e.m. **d-**Topographical distribution of the 25Hz steady state visual evoked potential amplitude as a grand average between patients and control subjects, for trials with 75 or 100% contrast foreground embedded in a midgray surround (0% contrast). The cluster of electrodes chosen to measure and statistically test ssVEP amplitude are marked with star symbols.

**Figure 4.**
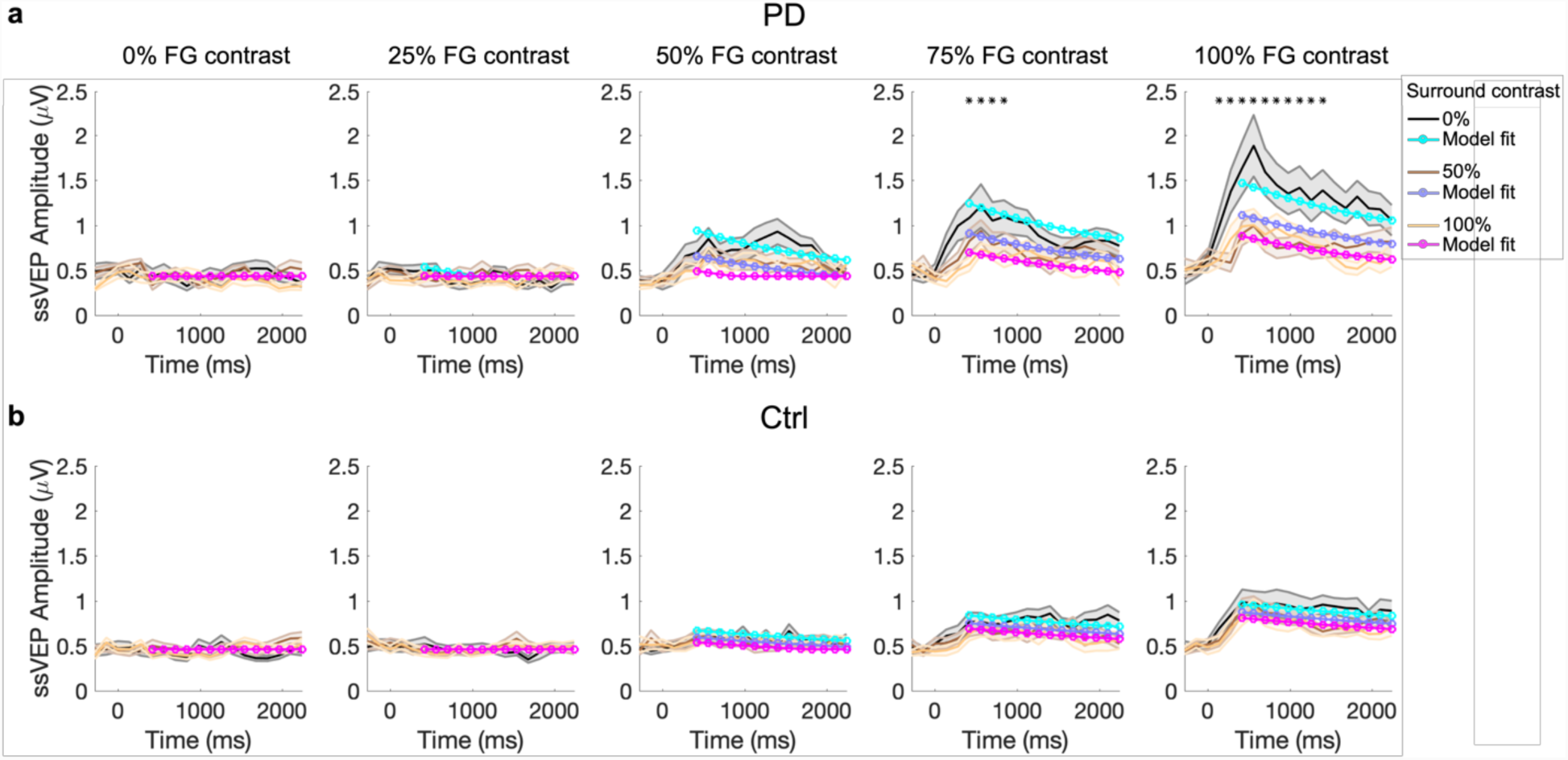
Temporal analysis. **A-** Patients with PD**. B-**Control subjects. The group average ssVEP amplitude was tracked over the duration of the visual stimulation, with stimulus onset at 0ms. We computed a short-time Fourier transform (STFT) on sliding windows of 560ms with an overlap of 75% over the 2.4-s trial length, starting from 280ms prior to stimulus onset. Temporal response profiles for increasing FG contrasts are plotted from left to right (0, 25, 50, 75, 100%) and each of the three surround contrasts are superimposed in each plot (black for 0%, brown for 50%, yellow for 100% contrast). The colored lines in cyan, purple and magenta represent the model fit to the data, in which FG and divisive surround drives each exponentially decay (i.e., adapt) to varying degrees. Shaded error bars indicate mean±s.e.m. Asterisks indicate timepoints with significantly higher amplitude of the visual response (ssVEP) in patients, reached for foreground stimulus of 75% and 100% contrast with surround contrast of 0% (p<0.05 in bootstrapping analysis).

As in our study of younger healthy subjects ^27^, we computed a summary metric of the surround suppression effect as the ratio between ssVEP amplitude corresponding to FG=100%, SS=100% and the ssVEP amplitude for FG=100%, SS=0% (Figure 3C). Suppression ratios were 0.46±0.04 for PD and 0.71±0.10 (mean±s.e.m) for control subjects, revealing a significantly stronger surround suppression effect in PD subjects (t_20_=-2.3304, p=0.0304). Thus, ssVEP amplitude showed a greater relative reduction in patients with PD than in controls under the presence of surrounding stimuli of increasing contrasts. We found no group difference in terms of contrast response function range (t_20_=0.8; p=0.43), computed as the difference between the ssVEP amplitude at maximum flicker contrast embedded in a midgray surround (FG=100%, SS=0%), and the ssVEP amplitude without either flicker or surround (FG=0%, SS=0%).

### Temporal response profiles

Temporal profiles of the visual response were constructed by plotting ssVEP amplitude over the duration of visual stimulation (short-time Fourier transform) for each stimulus condition. We found a steep increase of the visual response in the first 500ms after flicker onset, followed by a decay that reflected temporal adaptation. This was more evident for the high foreground contrasts. (Figure 4A,B). The results of an ANOVA showed in Table 1 (see Methods) revealed significant differences across groups that depended on contrast levels and time.

**Table 1.**
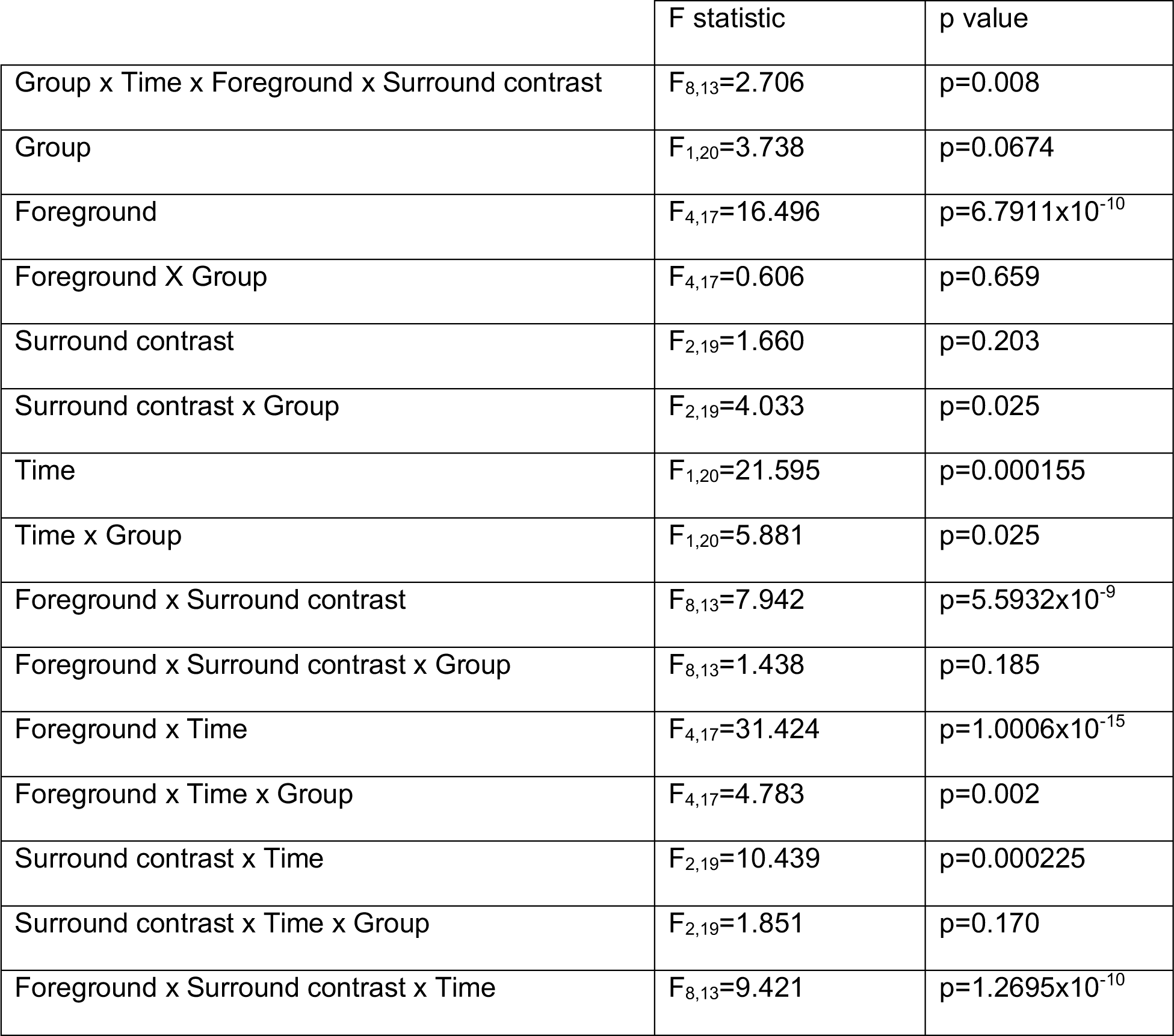
Factorial ANOVA showing the main effects and interactions between Group, Time, Foreground and Surround contrasts.

**Table 2.** Description of subject groups and clinical assessment scores in the final subject pool

|  | Parkinson's disease<br>patients (PD), n=11, 3<br>women | Healthy control subjects<br>(Ctrl), n=11, 7 women | t, p |
| --- | --- | --- | --- |
| Age (years) | 65.4 ± 4.5 | 66.7 ± 4.7 | $t_{20}=-0.69$, $p=0.498$ |
| MoCA | 27.6 ± 1.4 | 27.8 ± 1.7 | $t_{20}=-0.27$, $p=0.786$ |
| MHYS | 2.4 ± 0.3 | -- | -- |
| UPDRS Part<br>III | 26.8 ± 9.2 | -- | -- |

All values are given as mean ± standard deviation. An independent sample t-test was performed.

To unpack this, we conducted a bootstrapping analysis testing for group differences at each timepoint and for each contrast combination. We found significant differences only for time points between 420-840ms in the 75% FG, 0% SS condition, and for timepoints between 140-1400ms in the 100% FG, 0% SS condition (see asterisks, Figure 4). Since the amplitude increase in PD appeared to be restricted to earlier timepoints for the un-surrounded, high contrast stimuli, we compared with post-hoc test the rates of temporal decline. A Wilcoxon rank-sum test on the slope of a line fit over all timepoints from 500 to 1000 ms after stimulus onset indicated a significantly steeper, faster decay of the visual response in PD subjects (Z=-2.37; p=0.0179).

We next examined whether these differences could be described with a model with divisive normalization reflecting surround suppression and exponential decay functions to capture temporally adapting foreground and surround drives (see Methods). Shuffle statistics applied to the group-average data fits indicated a significant difference in suppression factor (β = 1.31 in PD and 0.338 in Controls), consistent with the above findings based on the suppression ratio. However, the difference in no other model parameters reached significance, including the time constant of temporal decay (τ) and the maximum amplitude Rm, as may have been expected from the results above.

Finally, we determined whether such visual abnormalities were related to characteristics of PD. We found no significant correlations between UPDRS motor scores and maximum ssVEP amplitude (ssVEP amplitude for FG=100%, SS=0% in the contrast response function), or suppression ratio (Pearson r=-0.11; p=0.759 and r=0.32, p=0.37, respectively). Similarly, disease duration was not significantly correlated to maximum ssVEP amplitude and suppression ratio (r=0.32; p=0.364, and r=0.53, p=0.116, respectively). Moreover, therapy did not significantly influence VEP measurements: in fact, the levels of levodopa equivalent dose did not correlate either with maximum ssVEP amplitude (r=0.101; p=0.780) or suppression ratio (r=0.057, p=0.875).

## Discussion

In this study, we examined the dynamics of visual responses in PD across different foreground and surround stimulus contrasts. Our results revealed three main characteristics of the visual response in PD patients and provide a new perspective on the visual deficits reported in this disease. First, in PD there was a greater visual response than in controls for flickering stimuli presented in a no-surround configuration: the visual response in patients peaked at a significantly higher level, and rapidly decayed over time. Second, patients exhibited a marked suppression of the visual response in the presence of surrounding stimuli. The influence of stimulus context upon the ssVEP amplitude was robustly expressed on the significant decrease of ssVEP amplitude embedded in a surrounding stimulus of 100%, the maximum contrast. Third, the EEG spectrum in PD showed a significantly higher level of background activity, or noise. These three findings challenge the notion that reduced visual contrast sensitivity and visual deficits observed in PD mainly stem from retinal dysfunction.

The higher spectral power in patients over the theta, alpha and beta bands (Figure 1) derived from occipital electrodes are in line with previous reports of increased spectral background over centroparietal regions in alpha and beta bands, mostly during dopaminergic therapy.^31-34^

The visual response in PD reached a higher initial amplitude and more quickly decayed for visual stimuli of 75 and 100% contrast and no surround. These results are in line with those of recent animal work. For instance, in non-human primates, modulation of dopaminergic forebrain circuits with D1 receptor antagonist injected in a prefrontal cortical area enhanced the visual response magnitude and orientation selectivity.^35^ Moreover, a PD model in drosophila exhibited similar sensory enhancement, reflected in response gain of the contrast response function.^36^ While the present findings are in agreement with such an enhancement of visual responses in PD, previous human studies in PD found in some cases, decreased VEP amplitudes or, in most cases, no amplitude changes at all.^8,37,38^ Differences in stimulus characteristics may explain such discrepancy: stimuli were usually presented in the center of the screen, covering a screen area from at least 4 to 18 degrees of visual angle or full screen, at 2 reversals per second.^8,13-16^ Due to retinotopic organization, evoked responses vary dramatically with stimulus location and size and for large foveal stimuli electric fields generated in V1 tend to self-cancel due to cortical folding.^28,39^ Since we have found an exaggerated degree of suppression in PD, one could speculate that with much larger stimuli, the greater mutual suppression within the stimulus cancels out any increase in bottom-up stimulus drive. A more likely explanation, however, may lie in the differences in spatial frequencies used – we avoid those previously found to illuminate retinally-mediated differences. In general, our stimulation paradigm offers a view on unique aspects of visual responses; whereas in previous work transient VEP responses were evoked by sequentially presented stimuli in 200-500 ms epochs, our steady-state paradigm enables tracing response dynamics continuously over a longer stimulation interval (2.4 s), revealing a characteristic temporal profile of rise and decay.

While both groups showed a decrease of ssVEP amplitude under the presence of surrounding stimuli, this surround suppression effect was proportionally stronger in patients. Visual surround suppression is known to be generated by a variety of lateral, feedforward and feedback connectivity across man hierarchical levels from retina ganglion and geniculate cells to striate and extrastriate areas.^29,30^ Indeed, in PD, degeneration of amacrine dopaminergic neurons might cause weakening of pre-cortical center-surround interactions in the retina. However, we found a significantly greater surround suppression effect in the cortical responses of PD patients as well as steeper temporal adaptation. Thus, we speculate that compensatory mechanisms arising at the cortical level may account for the lack of center surround interactions at the retinal level, and feedback from extrastriate areas to V1 may play a major role in such compensatory mechanisms,^29,30,40^ as it is also suggested by recent imaging studies.^41^

In our previous work, we found a relationship between the degree of electrophysiological surround suppression and perception.^27^ In particular, the reduction of the point of subjective equality of a stimulus when embedded in a high contrast surround, as demonstrated earlier ^42^, correlates with the decrease on the ssVEP amplitude measured using our paradigm. This suggests that in patients with PD, perception of a stimulus may be excessively decreased under the presence of a surround, a prediction that will require future testing. The exaggerated spatial and temporal gain control indices, coupled with the elevated level of noise in which visual responses are embedded, may plausibly contribute to the reduced visual contrast sensitivity often observed in PD patients. However, the spatial frequency of our stimuli was outside of the range previously found to exhibit deficits in PD^18,43^, and whereas contrast sensitivity effects are measured for low-contrast stimuli close to visual detection threshold, our effects were observed for much higher contrasts. Altogether, these findings highlight the need for systematic examination of a broader range of visual stimulus features and phenomena to attain a full understanding of visual system abnormalities in PD.

Electrophysiological indices of visual surround suppression may offer valuable insight into aspects of motor impairment in PD. For instance, the results of our study can explain why patients with PD and freezing of gait experience fewer freezing episodes while walking when visual cues are present on the floor.^44,45^ Indeed, salient stimuli help alleviate the freezing of gait episodes: one can thus speculate that the corridor resembles the “foreground” or central stimulus, embedded in “surrounding” objects, such as the doorway and walls. In PD, the pronounced surround suppression effect is hypothetically present in a natural environment, with strong contextual interactions that “suppress” responses to central stimuli. Since only salient visual cues like the lines provide a benefit to freezing, such cues may have the potential to overcome existing suppression of context upon a central stimulus. Of course, for now this remains a speculation to be tested; our study had an insufficient number (n=3) of patients with freezing to verify whether contextual effects were stronger in this subset of patients.

This study has some limitations. Indeed, perceptual effects were not assessed because of time constraints. Patients on chronic dopaminergic therapy were tested during regular medications schedule and thus, possible effects of chronic and acute treatment cannot be excluded. Also, the relatively small sample of patients warrants larger studies to confirm and extend the present findings. Finally, besides addressing these issues and the interaction between visual and motor system, future studies could couple suppression metrics with structural measurements in the retina that are linked to the visual dysfunction in PD.^46^

## Materials and Methods

### Subjects

EEG data were collected on the surround suppression paradigm as part of a larger multi-test study of 28 patients with PD (7 women) and 30 healthy control subjects (20 women), which adopted inclusive recruitment criteria that only required subjects to be right-handed and to show no signs of dementia. As described below, all subjects also received a complete neuropsychological assessment. Subjects were examined by a movement disorder neurologist using the standardized rating scales for cognitive and motor dysfunction: the Unified Parkinson Disease Rating Scale, UPDRS, and the Modified Hoehn and Yahr scales. All EEG recordings were conducted while patients were on their usual dopaminergic medication schedule. All subjects had normal or corrected-to-normal vision.

The protocol was approved by the Institutional Review Board of The City College of New York and NYU Langone School of Medicine. All participants signed an informed consent form before the test.

In the present study our final analysis was conducted on a subset of age-matched, cognitively unimpaired PD patients and control subjects who had sufficient trials for unbiased analysis of visual response magnitudes. Specifically, 10 controls and 10 PD patients were excluded for having MoCA scores of less than 26^47^ and/or a diagnosis of depression, and a further 4 controls and 7 PD patients were excluded on the basis that they did not complete a sufficient number of artifact-free trials to measure response amplitudes unconfounded by differences in background spectral EEG levels (see analysis details below). The final 11 control subjects were selected from the remaining 16 so that the two groups of 11 were matched for age and gender, with selections made randomly wherever multiple options for inclusion existed.

All values are given as mean ± standard deviation. An independent sample t-test was performed.

The AD8 Dementia Screening Interview, the 4 item mini physical performance test (MiniPPT) and the Hospital Anxiety And Depression Scale (HADS) revealed no signs of clinical depression or anxiety in the final pool of subjects, with no significant differences between the two subject pools (AD8: PD=1.55 ± 1.63, Ctrl=1 ± 1.61; t_20_=0.79, p=0.44; HADS Anxiety Scale: PD=4.27 ± 2.41, Ctrl=3 ± 2.32; t_20_=1.26; p=0.22; HADS Depression Scale: PD=3.64 ± 2.01, Ctrl=2.36 ± 2.06; t_20_=1.46; p=0.16; MiniPPT: PD=13 ± 1.67, Ctrl=14.1 ± 1.3; t_20_=-1.7; p=0.103).

### Experimental setup

Recordings were performed in a dark, sound-proof, radio-frequency interference-shielded room. Stimuli were presented dichoptically on a gamma-corrected CRT monitor (Dell M782) with a refresh rate of 100Hz and 800×600 resolution, at a viewing distance of 57cm. Stimulus presentation was programmed in MATLAB (6.1, The MathWorks Inc., Natick, MA, 2000), using the Psychophysics Toolbox extension.^48^

Surround suppression effects were tested using steady-state visual responses (ssVEP) to flickering foreground stimuli embedded in static surrounds, where stimuli in the upper field were flickered out-of-phase relative to those in the lower field. This method exploits anatomical and signal-summation principles to produce robust ssVEPs.^28^ The central “foreground” stimulus (FG) was composed of four vertically-oriented circular gratings, located at an eccentricity of 5 degrees of visual angle, at polar angles of 20° above (two upper disks) and 45° below (two lower disks) the horizontal meridian. Disks flickered on and off at 25 Hz, embedded within a non-flickering full-screen static “surround” (SS) also with vertical orientation, parallel to the FG. In this configuration, the “foreground” flickering stimulus is embedded into a “surrounding” pattern in order to induce maximum visual surround suppression effects using peripheral flickering foreground stimuli.^27^ In all cases, foreground and surround patterns were sinusoidally modulated luminance gratings with a spatial frequency of 1 cycle per degree. The average luminance of all gratings was equal to that of the midgray, 0% contrast surround, which was 65 cd/m^2^. Foreground contrasts could be 0% (midgray), 25%, 50%, 75% or 100% (black and white stripes) and static surround stimulus contrast could be 0%, 50% or 100%, both randomly assigned on each trial (see Figure 3 insets, stimulus configurations for FG 100% embedded in SS of 0% and 50% contrast).

For each of the contrast conditions, surround stimuli were presented spatially in-phase and opposite-phase relative to the foreground, interleaved pseudorandomly, and we collapsed across spatial phase conditions within each configuration to reduce border effects (see Vanegas, et al. ^27^). The full paradigm included a total of 120 trials (15 contrast combinations, 4 repetitions each for surround stimuli in and out of phase with foreground). Each trial started with the presentation of a fixation spot for 500ms, after which the flickering foreground and static surround stimuli were simultaneously presented for 2,400ms. Subjects were instructed to maintain their fixation on the center of the screen throughout each trial.

### EEG recordings

During each block, high-density electroencephalography data (HD EEG, Electrical Geodesics Inc., Eugene, OR) were recorded from 256 electrodes at a sample rate of 1,000Hz. We applied an online notch filter at 60Hz. Impedances were stable below 50kO. We selected 183 channels on the scalp and downsampled to 500Hz. Eye gaze was monitored continually using an EyeLink1000 (SR-Research) eye tracker to ensure gaze was within 1 degree of visual angle around the fixation spot during stimulus presentation.

### Data analysis

Analysis was carried out offline using in-house Matlab scripts in conjunction with topographic mapping functions of EEGLAB, an open source toolbox for EEG analysis.^49^ Individual epochs were extracted in the interval [−600:2600]ms relative to the visual stimulus onset at 0ms. Blinks were detected using an in-house blink detection script based on the frontal electrodes with a threshold of 60μV. Trial epochs were rejected if they contained a blink or an artifact greater than 300μV within the 2240-ms window beginning 160ms after stimulus onset, avoiding most of the early component of the transient visual evoked response. We baseline-corrected each trial epoch by subtracting the mean value over the points [−50:0]ms and re-referenced all channels to average mastoids by simple subtraction. We used Fourier decomposition to examine frequency spectra and compute contrast response functions from ssVEP amplitude corresponding to increasing levels of “foreground” contrast with and without “surrounding” patterns. The paradigm was designed to elicit visual sensory responses at a high frequency (25Hz), clear of the strongest endogenous rhythms such as alpha (10Hz). However, spectral amplitudes at any frequency bin are influenced by differences in levels of background noise. In PD, spectral power is known to be increased compared to controls.^31,33^ The power increase is mainly evident over centro-parietal regions in the alpha and beta ranges in levodopa-treated patients.^34^ We therefore began our analysis by examining the full EEG spectrum in the entire cohort of subjects by computing the Fast Fourier Transform (FFT) for the blank-screen (baseline) condition of FG=0% and SS=0% contrast (Figure 1 inset). We computed the FFT for a 2240-ms window beginning 160ms after each stimulus onset and then averaged across trials, and across electrodes over the central posterior midline region: Pz, POz and Oz, and neighboring electrodes, just as in Vanegas et al.^27^ This analysis using the entire subject cohort confirmed the background spectral difference in PD subjects as has been previously reported in the literature (Figure 1).^31,33^

This difference in background EEG noise levels could potentially confound comparisons of ssVEP amplitude between the two groups because it causes all frequencies, including the precise frequency of the ssVEP, to be elevated in PD. We thus carried out a noise-equalization procedure to preclude this confound, whereby we averaged the ssVEP measurements across fewer trials in the control subjects than in the PD patients in such a way that background spectral levels were equalized. Since background brain rhythms are asynchronous to the flickering onset, the background spectrum (noise) is expected to decrease with trial number when the FFT is computed on the average across trials. To determine the trial numbers required for noise-matching, we computed the frequency spectra for the blank-screen (0% foreground and 0% surround) condition as a function of number of trials. For each number of trials, we randomly selected this number 1,000 times from the available trials for this blank-screen condition and then averaged the results. We found that spectral background levels matched when the FFT was computed using the maximum trial number in PD and approximately half the available number of trials per condition in controls (Figure 2). We consequently applied this randomized trial selection, time-domain averaging and FFT procedure to the estimation of ssVEP amplitude in all task conditions. Since ssVEPs are locked to the flickering stimuli, signal averaging in the time domain reduces the background noise without affecting the amplitude of the average of the underlying oscillation. In this way, we equalized baseline noise levels across groups, so that any difference on the ssVEP was independent from the background spectral levels and not driven by differences in spectral profiles. Contrast response functions were derived by taking the number of trials for which equal levels of background spectral noise are achieved in PD and Ctrl groups.

Subjects were excluded if the minimum amount of trials per condition (7 to 8 for patients with PD and 4 for controls) was not met for each of the 15 contrast combinations after artifact rejection and blink detection. In the 11 PD and 11 Control subjects included in the final analysis, we verified that the levels of background noise were still equalized using a t-test (t_20_=0.689, p=0.498).

### Contrast response functions

In order to examine the influence of contextual surrounding patterns on the ssVEP, we derived contrast response functions as described in Vanegas, et al.^27^ A fast Fourier transform (FFT) was computed for a 2,240-ms window beginning 160ms after stimulus onset to extract the single amplitude value at the frequency of stimulation 25Hz. To examine temporal dynamics, including the timecourse of adaptation, we computed a short-time Fourier transform (STFT) on sliding windows of 560ms with an overlap of 75% over the 2.4-s trial length, starting from 280ms prior to stimulus onset (Figure 5).

**Figure 5.**
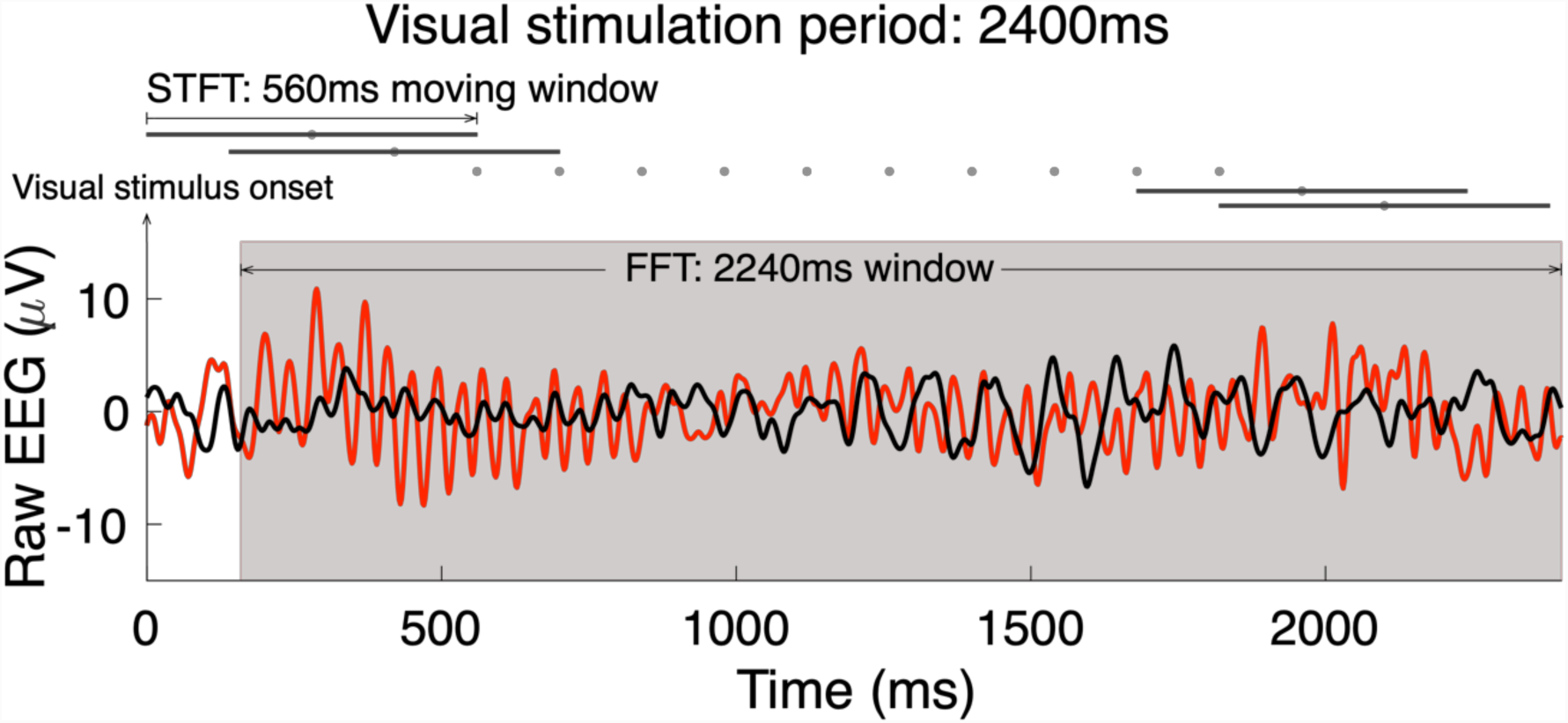
Methods of analysis of the visual response. Example raw EEG traces in a PD patient (red) and a Ctrl subject (black), measured from a cluster of parieto-occipital electrodes during a visual stimulation trial, FG=100% and SS=0% contrast. Time frequency analyses, FFT and STFT, were performed to extract the ssVEP amplitude at the frequency of stimulation 25Hz.

### Model fit

As in our original study using this paradigm in young healthy subjects,^27^ we fitted the grand-average time-resolved contrast response functions with a model accounting for variations over time as well as foreground and surround contrast. First, the basic amplitude increase as a function of foreground contrast was captured by the standard Naka-Rushton function.^50^ Second, the influence of the spatial surround was encapsulated in an additive term in the denominator. Third, the temporal adaptation of both the foreground and surround drives was described by decaying exponentials with the same time constant across contrasts levels and stimuli (foreground and surround) but asymptotes that were permitted to differ. The complete model is described by the relation:

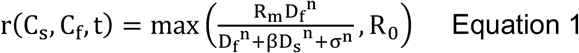

where r is the ssVEP amplitude at time t, R_m_ is the maximal response, R_0_ is the spectral baseline noise, σ is the contrast at which half of the maximum response is achieved, β is the coefficient of suppression, which scales the influence of surround contrast in the denominator and n is the exponent that accounts for non-linearity of the function. Since in spectral EEG measurements the baseline level R_0_ reflects the noise floor (25-Hz amplitude even in the absence of any stimulation), we use the max function to reflect the fact that measurements cannot pass below this noise floor. The time-dependent foreground drive D_f_(t) and suppressive drive D_s_(t) were modeled as decaying exponential functions beginning at the veridical physical contrast of the stimulus and asymptotically tending towards a fraction of that value, captured in the relations:

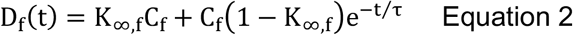

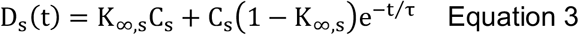

where τ is the mutual time constant for adaptation of foreground and surround drives and C_f_ and C_s_ are the foreground and surround contrasts, respectively. K_∞,f_ and K_∞,s_ represent factors by which the foreground and suppressive drive are asymptotically reduced relative to the initial value. The fit was carried out on the data over the interval 420 to 2240ms so that FFT windows stayed within the bounds of the stimulation period and using the method of least-squares. The best fit parameters of the model (see equations 1, 2 and 3) were as follows: PD subjects: R_m_=2.94, R_0_=0.43, n=1.08, σ=1, β=1.31, τ=1371.15, K_∞,f_=0.44, K_∞,s_=0.77; Ctrl subjects: R_m_=1.92, R_0_=0.46, n=0.9, σ=1, β=0.34, τ=2549.21, K_∞,f_=0.49, K_∞,f_=1.

### Statistical analyses

To first of all test for differences in ssVEP amplitude as a function of group, time, foreground contrast and surround contrast, and/or interactions between them, we carried out a mixed 4-way ANOVA with two time points taken early (420-560ms) and late (1540-1680ms) in the stimulation epoch. To follow-up the interactions found in this test we assessed ssVEP amplitude differences across groups for all individual conditions using shuffle statistics with 1,000 sampling iterations (random reassignment of group). Shuffling was done per time point over the duration of the visual stimulation (short-time Fourier transform), per stimulus condition. To test for spectral differences in the blank-screen condition, we performed a bootstrap analysis at each frequency bin. In this method, statistical samples were obtained by resampling from the shuffled original sample 1,000 times. Surround suppression effects and clinical scores were also tested using independent samples t-tests in cases where measures met the Kolmogorov-Smirnov test for normality. When samples were not normally distributed (i.e., slope of the line, temporal response), we conducted a nonparametric Wilcoxon rank-sum test for independent groups. Correlation analyses (Pearson) were computed to test for relationships between two clinical scores (i.e., duration of the disease and motor score in the UPDRS scale, in patients) and two electrophysiological metrics: 1-ssVEP amplitude in the stimulus condition of FG=100% and SS=0%, and 2-surround suppression effect or suppression ratio, calculated as the ratio between ssVEP amplitude corresponding to FG=100%, SS=100% and the ssVEP amplitude for FG=100%, SS=0%. In order to test for significant differences between the model parameter values fit to the grand average of each group, we used shuffle statistics; specifically, we randomly reassigned group membership 500 times and fit the model to the two false ‘groups’ each time to form a null distribution of parameter value differences across groups, and compared the true parameter ‘value’ difference against this null distribution. The level of significance was set at 0.05 in all tests.

## Acknowledgements

Research reported in this publication was supported by the Michael J Fox Foundation, the Safra Foundation, the National Parkinson Foundation (to MFG and ADR), the National Institutes of Health (R01 NS054864 to MFG), grants from the National Institute of General Medical Sciences (SC2-GM-099626 to SPK) and from the National Science Foundation (BCS-1358955 to SPK).

## Data availability

The datasets generated and analyzed in this study are available from the corresponding authors on reasonable request.

## Author contributions

M.I.V., M.F.G and S.P.K. designed the experiments. M.I.V. and A.B. conducted the experiments. M.I.V. and S.P.K. analyzed the data. M.I.V., A.B., J.G., A.D.R., A.Q., M.F.G. and S.P.K. wrote the paper.

